# A wave of minor *de novo* DNA methylation initiates in mouse 8-cell embryos and co-regulates imprinted X- chromosome inactivation with H3K27me3

**DOI:** 10.1101/2023.10.06.561284

**Authors:** Yuan Yue, Wei Fu, Qianying Yang, Chao Zhang, Wenjuan Wang, Meiqiang Chu, Qingji Lyu, Yawen Tang, Jian Cui, Xiaodong Wang, Zhenni Zhang, Jianhui Tian, Lei An

## Abstract

DNA methylation is extensively reprogrammed during early stage of mammalian development and is essential for normal embryogenesis. It is well established that mouse embryos acquire genome-wide DNA methylation during implantation, referred to as *de novo* DNA methylation, from globally hypomethylated blastocysts. However, the fact that the main *de novo* DNA methyltransferase 3B (DNMT3B) is initially expressed as early as the 8-cell stage, contradicts the current knowledge about timing of initiation of *de novo* DNA methylation. Here, we reported that a previously overlooked minor wave of *de novo* DNA methylation initially occurs during the transition from the 8-cell to blastocyst stage, before the well-known large-scale *de novo* DNA methylation during implantation. Functional analyses indicated that minor *de novo* DNA methylation regulates proliferation, lineage differentiation and metabolic homeostasis of preimplantation embryos, and is critical for embryonic developmental potential and pregnancy outcomes. Furthermore, bioinformatic and functional analyses indicated that minor *de novo* DNA methylation preferentially occurs on the X chromosome and co-regulates imprinted X-chromosome inactivation via the interaction between DNMT3B and polycomb repressive complexes 2 core components during blastocyst formation. Thus, our study updates the current knowledge of embryonic *de novo* DNA methylation, thereby providing a novel insight of early embryonic epigenetic reprogramming.

**Summary statement:** A minor wave of *de novo* DNA methylation has been initiated prior to blastocyst formation, but not during the implantation period, and co-regulates imprinted X-chromosome inactivation.

## INTRODUCTION

Mammalian early embryos are co-regulated by a series of spatiotemporally restricted epigenomic events that are critical for acquiring developmental potential. In mouse, it has been widely accepted that a wave of genome-wide *de novo* DNA methylation occurs around the time of implantation, thus re-establishing DNA methylation patterns from a globally hypomethylated blastocysts to relatively static DNA methylation patterns in post-implantation embryos (Borgel et al., 2010; Smith et al., 2012a). DNA methyltransferase 3B (DNMT3B) is the main enzyme that catalyzes embryonic *de novo* DNA methylation. Genetic depletion of *Dnmt3b* results in hypomethylation and dysregulation of pluripotency genes, gastrulation genes, germline-specific genes, *etc* (Auclair et al., 2014; Borgel et al., 2010), thus causing severe developmental defects and lethality after implantation (Li et al., 1992; Okano et al., 1999). However, despite its developmental importance, the exact timing of initiation of *de novo* DNA methylation remains largely elusive.

Although earlier results using high-throughput sequencing methods suggested that *de novo* DNA methylation occurs during the transition from blastocysts to early post-implantation embryos (Auclair et al., 2014; Borgel et al., 2010; Guo et al., 2014; Smith et al., 2012a), our detection assays (Figure 1A, B) and other previous studies (Guenatri et al., 2013; Hirasawa et al., 2008; Li et al., 2016) indicated that *Dnmt3b,* the main *de novo* DNA methyltransferase, is initially expressed as early as the 4-cell stage, and shows clear nuclear localization at 8-cell stage. This fact challenges current knowledge about timing of initiation of embryonic *de novo* DNA methylation, and therefore raising the possibility that embryonic *de novo* DNA methylation may precede, but not follows, the blastocyst stage.

**Figure 1.**
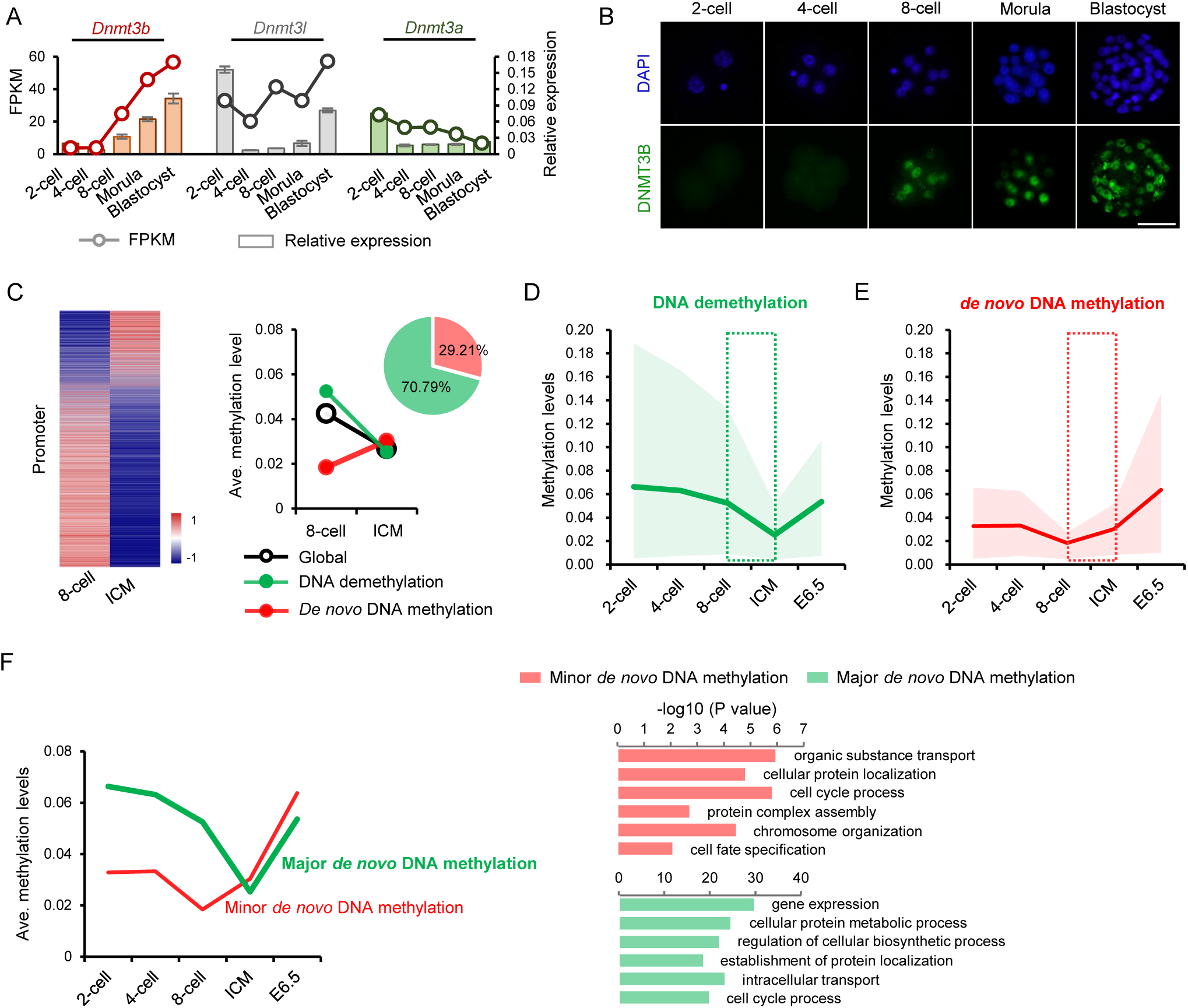
A small proportion of promoters initiate *de novo* DNA methylation by 8-cell stage. (A) The mRNA expression levels of *Dnmt3s* from the 2-cell to blastocyst stage. The left ordinate represents FPKM values of *Dnmt3s* from published transcriptome data (Fan et al., 2015), and the right ordinate represents relative expression levels detected by qRT–PCR. (B) Immunofluorescent staining of DNMT3B in mouse preimplantation embryos. Scale bars: 50 µm. (C) DNA methylation changes of promoters during the transition from the 8-cell to blastocyst stage. Left panel: heat map of DNA methylation levels at promoters. Right panel: general trends of promoters undergoing *de novo* methylation or demethylation, and their relative contribution to differentially methylated regions. (D-E) DNA methylation dynamics of promoters subject to demethylation (D) or *de novo* methylation (E) during the transition from the 8-cell to blastocyst stage (red and green lines, average methylation levels; colored space, 10th/ 90th percentile). (F) A model illustrating two waves of *de novo* DNA methylation before and after the blastocyst stage, and their potential functions.

To test this possibility, we reanalyzed well-established bisulfite sequencing data that profiles DNA methylation maps from the mouse preimplantation to post-implantation embryos (Smith et al., 2012a), and found approximately one-third of genes gained promoter DNA methylation during the transition from the 8-cell to blastocyst stage. It means that despite global DNA demethylation throughout the preimplantation development, a small proportion of regions have initiated *de novo* DNA methylation prior to the blastocyst stage, which we designated as minor *de novo* DNA methylation. We next ask its developmental functions for embryogenesis. The results showed that minor *de novo* methylation also regulates proliferation, lineage differentiation and metabolism of preimplantation embryos, and thus affecting the developmental potential and final pregnancy outcomes. Of interest, minor *de novo* DNA methylation is more prone to occur on the X chromosome. We next confirmed that minor *de novo* DNA methylation plays an important role in establishing imprinted X chromosome inactivation (iXCI) via the interaction between DNMT3B and the polycomb repressive complexes 2 (PRC2) core components, which are critical for X-linked heterochromatin. Thus, our study provides a novel understanding of *de novo* DNA methylation during early development, which updates the current knowledge of embryonic epigenomic hallmark events and their interactions.

## RESULTS AND DISCUSSION

### A small proportion of promoters initiate *de novo* DNA methylation by the 8-cell stage

To investigate the earliest initiation of *de novo* DNA methylation, we first examined the expression patterns of *Dnmt3s* in mouse preimplantation embryos. Results from previously published transcriptome (Fan et al., 2015) and our quantitative RT-PCR (qRT-PCR) detection showed that expression of *Dnmt3b*, the main enzyme for embryonic *de novo* DNA methylation (Borgel et al., 2010; Okano et al., 1999), as well as its non-catalytic partner *Dnmt3l* (Gowher et al., 2005), initiated by the 8-cell stage, and were further increased upon preimplantation development (Figure 1A). By contrast, the expression of *Dnmt3a*, which is mainly required for gametic methylation (Kaneda et al., 2004a), was maintained at low levels concurrently (Figure 1A). The initial expression of *Dnmt3b* was also confirmed at the protein level. Nuclear location of DNMT3B can be clearly detected from the 8-cell stage onward (Figure 1B). Given the essential role of DNMT3B in catalyzing *de novo* DNA methylation, we postulated that embryonic *de novo* DNA methylation may initiate as early as the 8-cell stage, which challenges current knowledge that *de novo* DNA methylation initiates during implantation.

To test the hypothesis, we next reanalyzed a well-established reduced representation bisulfite sequencing (RRBS) data that profiles DNA methylation maps from the mouse preimplantation to post-implantation stage (Smith et al., 2012a). Promoter regions are important ’s target sites of DNMT3B (Choi et al., 2011). The acquisition of DNA methylation in promoters, especially in intermediate and low CpG promoters, during implantation is largely dependent on DNMT3B, and plays an important role in regulating developmental genes (Auclair et al., 2014; Borgel et al., 2010; Dahlet et al., 2020). Thus, among genomic regions that may undergo *de novo* DNA methylation, we initially focused our analysis on DNA methylation dynamics of promoters from the 8-cell stage onward. Despite the global loss of promoter DNA methylation (Figure 1—figure supplement 1A), more detailed characterization showed that a considerable proportion of promoters (approximately 30%) gained DNA methylation during the transition from the 8-cell to blastocyst stage, even though most promoters underwent demethylation (Figure 1C). Similar observation was also obtained by reanalyzing another published whole-genome bisulfite sequencing (WGBS) data (Smith et al., 2017) (Figure 1—figure supplement 1B). In addition, a reanalysis based on a publicly available single cell multi-omics sequencing (COOL-seq) data of mouse early embryos (Guo et al., 2017) showed that both male and female embryos gain DNA methylation during the transition from the 8-cell to blastocyst stage (Figure 1—figure supplement 1C, D).

Many previous analyses using different methodologies have showed the promoter regions are relatively hypomethylated in general compared with other genome elements (Auclair et al., 2014; Borgel et al., 2010; Dahlet et al., 2020; Smith et al., 2012b). Despite this, by tracking DNA methylation dynamics of two subsets of promoters that gained or lost DNA methylation from the 8-cell to blastocyst stage respectively, we found promoters that underwent demethylation showed a relatively higher DNA methylation levels following fertilization, and then were gradually demethylated until reaching their lowest value in inner cell mass (ICM) (Figure 1D). In contrast, promoters that gained DNA methylation from the 8-cell to blastocyst stage, showed a relatively lower DNA methylation levels, following fertilization, and reached their lowest DNA methylation levels at the 8-cell stage (Figure 1E). Based on the low basal levels, these promoters gain DNA methylation from the 8-cell to ICM, and then continued to gain DNA methylation during implantation (Figure 1E). To distinguish two waves of *de novo* DNA methylation before and after the blastocyst stage, we designated them as minor and major *de novo* DNA methylation, respectively. By performing gene ontology enrichment analysis of genes with promoter regions that undergo minor or major *de novo* DNA methylation respectively, we noticed that besides of many important basic processes common to two waves of *de novo* DNA methylation, genes subject to minor *de novo* DNA methylation have been enriched in processes such as organic substance transport, chromosome organization, and cell fate specification (Figure 1F). Compared with conventionally accepted major *de novo* DNA methylation, minor *de novo* DNA methylation showed a smaller increase in promoter DNA methylation that tended to occur in low CpG-containing promoters (Figure 1—figure supplement 1E).

### Minor *de novo* DNA methylation participates in embryonic proliferation, differentiation and affects developmental potential

It has been known that *de novo* DNA methylation during implantation stage plays an essential role in transcriptional regulation of developmental genes that are critical for embryonic differentiation and survival (Auclair et al., 2014; Borgel et al., 2010), thus we next attempted to test the short-and long-term developmental consequences of minor *de novo* DNA methylation. To this end, we performed an integrated analysis using the publicly available bisulfite sequencing data (Smith et al., 2012a) and transcriptome data (Wu et al., 2016) that profiles DNA methylation and gene expression maps from the 8-cell to ICM, respectively. Putative genes transcriptionally regulated by minor *de novo* DNA methylation (Figure 2—figure supplement 1A) were functionally related to many basic processes, such as cell differentiation and metabolic regulation. Phenotype annotations in Mouse Genome Informatics (MGI) showed that genes enriched in these basic processes are critical for normal embryonic survival, development and postnatal growth (Figure 2—figure supplement 1B, C). It should be noted that genes subject to DNA demethylation also participate in similar processes (Figure 2—figure supplement 1D). In line with our results, previous studies using *Tet1*-deficient or *Tet1*/*3-*deficient embryos, indicated the critical role of TET-mediated DNA demethylation in these processes, as well as embryonic and postnatal development (Dawlaty et al., 2011; Ito et al., 2010; Kang et al., 2015). Thus, it can be presumed that minor *de novo* DNA methylation, and TET-mediated demethylation, may synergistically co-regulate DNA methylation homeostasis before blastocyst formation, thus fine-tuning embryonic developmental potential and long-term outcomes, enriching the knowledge of developmental significance of DNA methylation homeostasis in early embryos, as proposed previously (Iurlaro et al., 2017; Smith and Meissner, 2013).

We next wanted to examine whether minor *de novo* methylation is functionally associated with these basic biological processes before blastocyst formation and in turn affects developmental potential and pregnancy outcomes. Given identification of long-term developmental consequence of minor *de novo* DNA methylation requires that only minor, but not major *de novo* methylation, is functionally inactivated before blastocyst formation, we knocked down *Dnmt3b* in preimplantation embryos by microinjecting *Dnmt3b* siRNA into zygotes (Figure 2—figure supplement 2A-E). As expected, *Dnmt3b* mRNA and protein levels were significantly reduced in morulae, and tended to be lower in blastocysts compared to those of the negative control (NC) group (Figure 2—figure supplement 2A-C). The inhibition of *Dnmt3b* resulted in a significant decrease in the global level of 5mC staining before blastocyst formation (Figure 2—figure supplement 2D, E). *Dnmt3b* deficiency not only inhibited proliferation (Figure 2B), but also changed lineage differentiation of blastocysts (Figure 2C). The decreased EdU-positive signals, as well as the slight lower total cell number, suggested an inhibited proliferation due to *Dnmt3b* knockdown (KD). Since energy metabolism is crucial for supporting embryo differentiation and development, and oxidative phosphorylation (OXPHOS) is activated during the blastocyst stage (Zhao et al., 2021), we next examined the energy metabolism, in particular OXPHOS activity, of *Dnmt3b*-KD embryos. Despite the unchanged total ATP production, *Dnmt3b* deficiency led to a significantly disruption in OXPHOS-dependent-ATP production (Figure 2D). These observations were further confirmed via the transient treatment of 5-aza-dC specifically during the window of minor *de novo* DNA methylation (Figure 2—figure supplement 3A-D). 5-aza-dC is a well-established global DNA hypomethylating agent that efficiently inhibit the activity of all DNMTs and thus has been frequently used to study the maintenance of DNA methylation and *de novo* DNA methylation, despite its inhibitory effect common to various DNMTs (Maslov et al., 2012; Oka et al., 2005). Transient treatment of embryos with 5-aza-dC specifically during the window of minor *de novo* DNA methylation allows us to test its function in a temporally specific manner. Thus, it can be presumed that the disrupted proliferation and energy metabolism, appears not to be due to defective major *de novo* DNA methylation. Finally, we transferred *Dnmt3b*-KD embryos into pseudopregnant recipient females. Compared with that in NC group, *Dnmt3b*-KD embryos resulted in a significantly reduced implantation rate and live birth rate (Figure 2E-F), suggesting the important role of minor *de novo* DNA methylation in influencing embryonic developmental potential.

**Figure 2.**
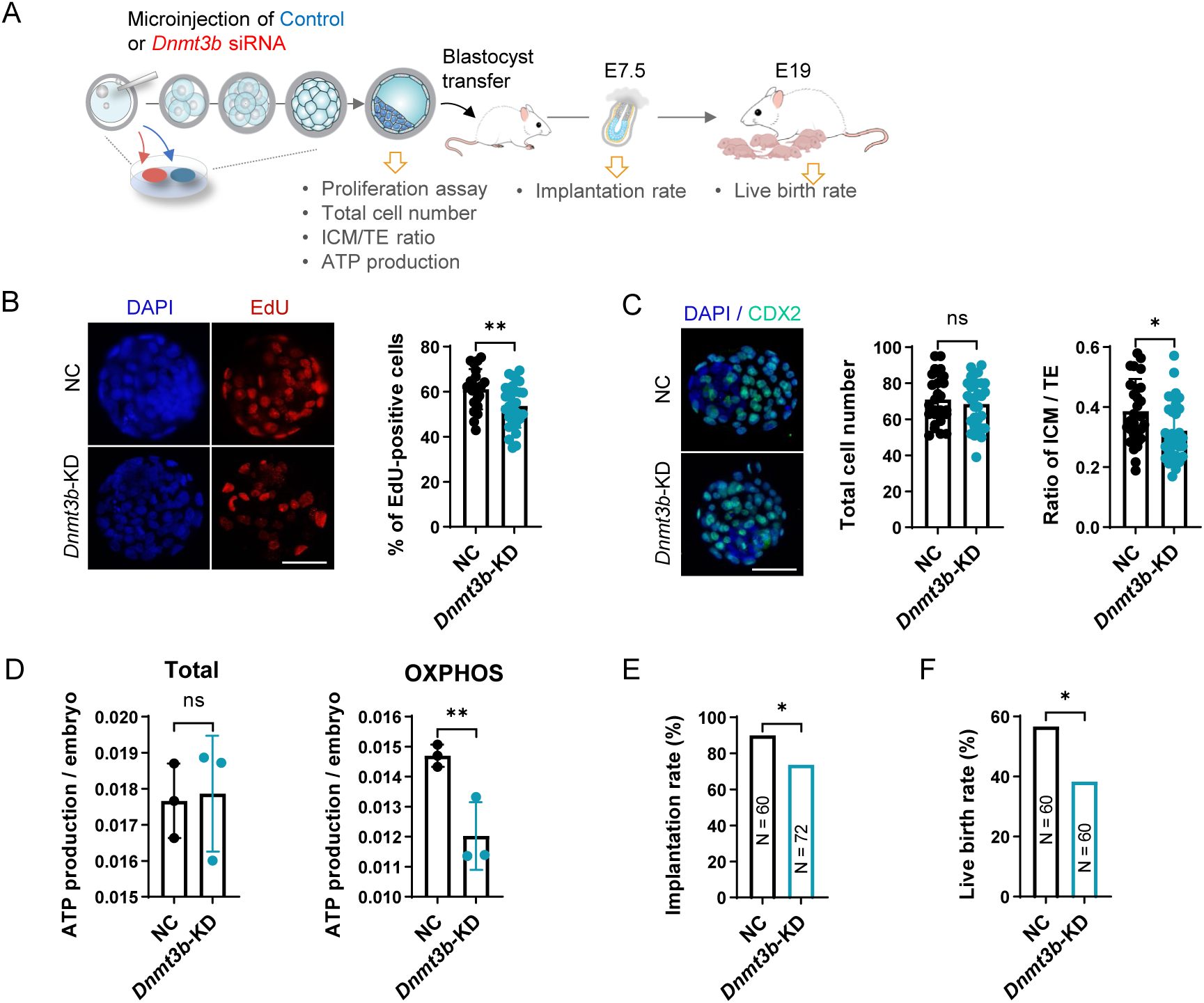
Minor *de novo* DNA methylation participates in embryonic proliferation, differentiation and affects developmental potential. (A) Schematic diagram of the experimental design. (B) EdU staining of NC and *Dnmt3b*-KD blastocysts. Right panel: the percentage of EdU-positive cells. (C) The total cell number (left panel) and the ratio of ICM/TE (right panel) in NC and *Dnmt3b*-KD blastocysts. (D) The total ATP production (left panel) and OXPHOS-dependent ATP production (right panel) in NC and *Dnmt3b*-KD blastocysts. (E-F) Implantation rate (E) and live birth rate (F) in NC and *Dnmt3b*-KD groups. All data are presented as the mean ± SD of at least three independent experiments. \**P* < 0.05, \*\**P* < 0.01. ns, not significant. Scale bars: 50 µm (A, B).

### Minor *de novo* DNA methylation preferentially occurs on the X chromosome and plays an important role in iXCI

To obtain a global view of minor *de novo* DNA methylation, we next detected the chromosome-wide distribution of *de novo* methylated promoters from the 8-cell to ICM, and found that they were globally distributed across autosomes, with the percentage ranging from 23% to 33% (Figure 3A-B). Of note, approximately half of X-linked promoters (48%) underwent minor *de novo* DNA methylation throughout the X chromosome (Figure 3—figure supplement 1A), much higher than those of autosomal promoters (Figure 3B). This result was similar with that of major *de novo* DNA methylation during implantation (Figure 3—figure supplement 1B, C). Using the single cell COOL-seq data, we also specifically reanalyzed the DNA methylation changes on the X chromosome in female embryos. The X chromosome showed a more notable increase than that on autosomes, and the female X chromosome showed a higher DNA methylation level than that of the male (Figure 3—figure supplement 2A, B). The X chromosome-biased prevalence of *de novo* DNA methylation is reminiscent of XCI, a female-specific epigenetic event that occurs during early development for balancing the X-linked gene dosage between males and females. In mice, early embryos undergo two waves of XCI: iXCI initiates and occurs specifically on the paternal X chromosome in preimplantation embryos, and then persists in extraembryonic tissues; while random X chromosome inactivation (rXCI) occurs randomly on either the maternal or paternal X chromosome in the embryonic cells by implantation stage (Augui et al., 2011; Chow and Heard, 2009; Lee and Bartolomei, 2013). Spatiotemporal co-occurrence of minor *de novo* DNA methylation and iXCI on the X chromosome, led us to ask if these two epigenetic events are functional linked during preimplantation development.

**Figure 3.**
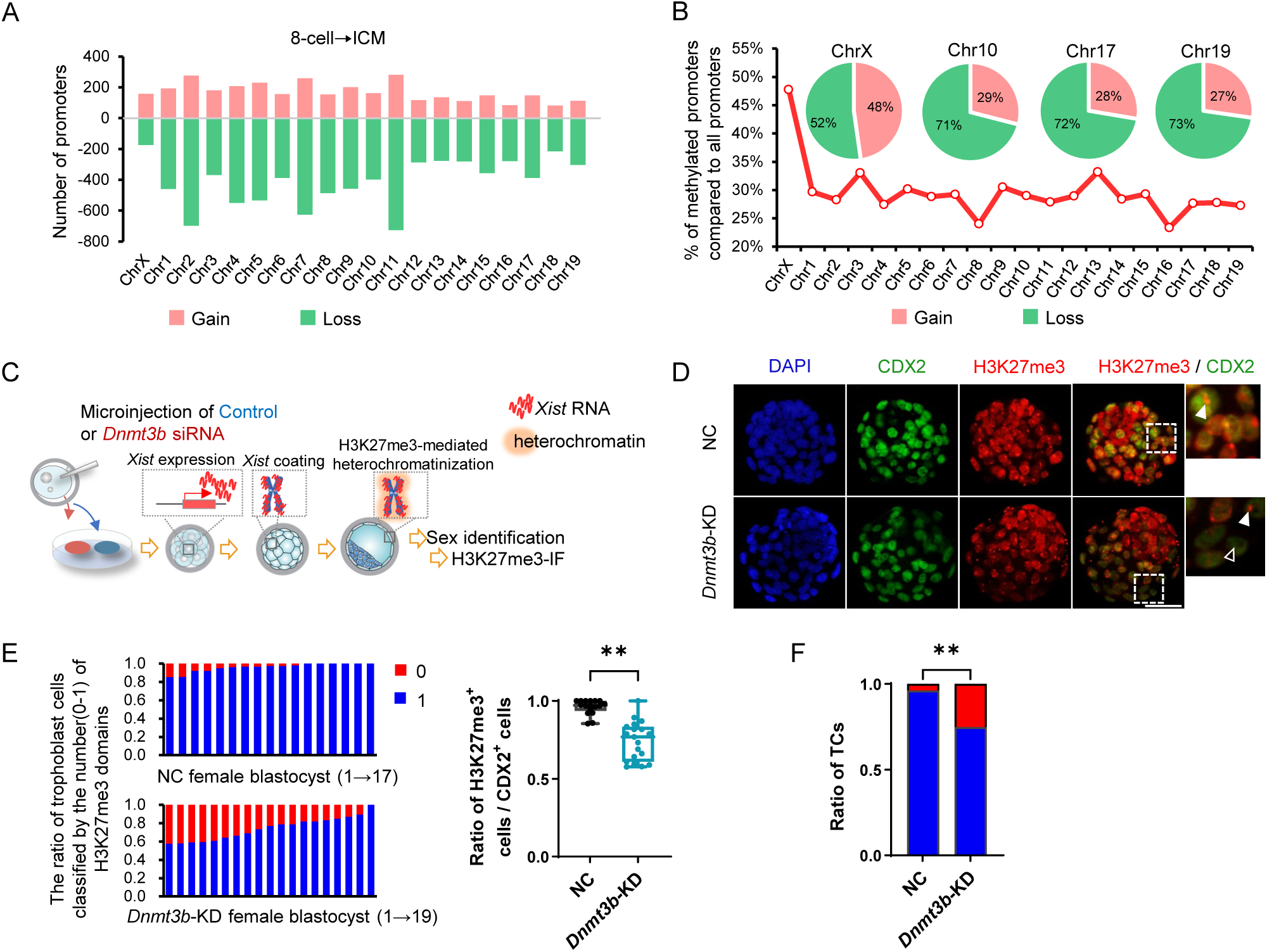
Minor *de novo* methylation preferentially occurs on the X chromosome and plays an important role in iXCI. (A) The chromosome-wide distribution of *de novo* methylated promoters during the transition from the 8-cell to blastocyst stage. (B) Percentage of *de novo* methylated promotors on each chromosome during the transition from the 8-cell to blastocyst stage. (C) Schematic diagram of the experimental design and the molecular processes of iXCI initiation and establishment. *Dnmt3b* knockdown was performed by microinjecting *Dnmt3b* siRNA into zygotes. Immunostaining assay for H3K27me3 was performed and quantified in trophoblast cells in each female blastocyst. (D) Representative immunostaining for H3K27me3 (red) in the nuclei (DAPI) of NC and *Dnmt3b*-KD female blastocysts colabeled with CDX2 (green)-positive trophoblast cells. Scale bars: 50 µm. The arrowhead indicates the H3k27me3 domain and the blank arrowhead indicates the blastomere without the H3k27me3 domain. (E) The ratio of trophoblast cells classified by the number of H3K27me3 domains in each NC and *Dnmt3b*-KD female blastocysts. Each bar represents one female embryo. (F) The percentage of H3K27me3-positive and H3K27me3-negative trophoblast cells (TCs) to total trophoblast cells in female blastocysts. All data are presented as the mean ± SD of at least three independent experiments. \*\**P* < 0.01. ns, not significant.

Earlier studies have focused on whether the DNA methylation imprinting established during oogenesis as the imprinting mark responsible for the monoallelic expression of *Xist* (the initiation event of iXCI) in preimplantation embryos and demonstrated that oocyte DNA methylation is dispensable for *Xist* imprinting (Chiba et al., 2008; Kaneda et al., 2004b; Sado et al., 2004). However, the role of DNA methylation in achieving X chromosomal heterochromatinization (the late event of XCI) remains controversial, although it has been well-accepted DNA methylation is tightly associated with maintaining X chromosome-wide transcriptional silence of rXCI during implantation (Blewitt et al., 2008; Sado et al., 2000). Recently, targeted DNA demethylation of genes on the X chromosome, using inactive Cas9 protein (dCas9) fused with the catalytic domain of ten-eleven translocation dioxygenase 1 (TET1), supported the causal role between DNA methylation and XCI status (Halmai et al., 2020). However, the conclusion is based on rXCI, the role of DNA methylation in the heterochromatinization of iXCI remains unclear, and this issue has been documented as a long-standing open question of epigenetic programming during preimplantation development (Csankovszki et al., 2001; Kaneda et al., 2004b; Sado et al., 2000; Sado et al., 2004). Actually, it is even plausibly thought that DNA methylation is not required for achieving heterochromatinization of iXCI because preimplantation embryos is undergoing global and massive DNA demethylation. To identify the biological function of minor *de novo* DNA methylation in iXCI, we knocked down *Dnmt3b* in preimplantation embryos by microinjecting *Dnmt3b* siRNA into zygotes (Figure 3C). Next, we detected the H3K27me3, a classic marker for establishment of XCI achieving X chromosome-wide heterochromatinization of transcriptional depression (Chow and Heard, 2009; Huynh and Lee, 2005; Wutz, 2011). In line with the decreased DNA methylation levels (Figure 2—figure supplement 2D, E), we found nearly all detected *Dnmt3b*-KD female blastocysts contained trophoblast cells lacking the H3K27me3 domain, with a variable frequency ranging from 10.7% to 42.4%; whereas nearly all NC female blastocysts showed a single H3K27me3 domain in each trophoblast (Figure 3D-E). The proportion H3K27me3-positive cells in total trophoblast cells were significantly lower due to *Dnmt3b*-KD (Figure 3F), indicating *Dnmt3b* deficiency leads to an impaired iXCI in preimplantation female embryos. This result was confirmed using *Dnmt3b*-knockout (KO) embryos (Figure 3—figure supplement 3A-D). To further validate the function of minor *de novo* DNA methylation in iXCI, we take advantage of chemical-induced DNMT inhibition using 5-aza-dC (Figure 3—figure supplement 3E-H). In addition, the involvement of minor *de novo* DNA methylation in iXCI establishment was also supported by the finding that many representative X-linked non-escaping genes tended to be *de novo* methylated during the transition from the 8-cell to ICM (Figure 3—figure supplement 3I-J).

To test whether DNA methylation loss leads to a prolonged effect on iXCI, we further reanalyzed the RNA-seq and H3K27me3 CHIP-seq data in extraembryonic ectoderm (ExE) of E6.5 single embryos that underwent *Dnm3a*/*3b* knockout because preimplantation iXCI status maintains in extraembryonic cells (Chen et al., 2019; Galupa and Heard, 2015; Schulz and Heard, 2013). The results showed that chromosome-wide loss of DNA methylation led to a nearly complete loss of H3K27me3 on paternal X chromosome (specifically inactivated in iXCI), along with a notable transcriptional upregulation cross the chromosome, including the locus of some non-escaping genes. By contrast, these changes cannot be not observed on maternal X chromosome (Figure 3—figure supplement 4A, B).

Taken together, these data suggest that a wave of minor *de novo* DNA methylation preceding the blastocyst stage, participate in the establishment of iXCI. Although a recent study reported that *Dnmt3b* has a dominant role on the X chromosome in mouse embryonic fibroblasts (Yagi et al., 2020), the detectable DNMT3A/*Dnmt3a* in preimplantation embryos, albeit at low levels (Uysal et al., 2017; Zhang et al., 2015), led us to speculate that DNMT3B and DNMT3A synergistically act on minor *de novo* DNA methylation, similar with that reported in subsequent peri-implantation embryos (Borgel et al., 2010; Chen et al., 2019).

### Minor *de novo* DNA methylation co-regulates iXCI via the interaction between DNMT3B and PRC2 core components

Having confirmed the functional role of minor *de novo* methylation in iXCI, we next explored the underlying mechanism. Because iXCI is initiated by *Xist* RNA upregulation and coating during the 8-cell and morular stage (Penny et al., 1996), we first detected *Xist* mRNA expression. No significant difference in the degrees of absence of the *Xist* domain in each blastomere nucleus between NC and *Dnmt3b*-KD female morulae (Figure 4A, B), which were also confirmed by detecting *Dnmt3b*-KO female morulae (Figure 4—figure supplement 1A, B). In line with this, single female embryo qRT-PCR analysis of *Xist* and its essential upstream activator *Rnf12* during initial stage of iXCI, showed that the expression levels of *Xist* and *Rnf12* were not affected by *Dnmt3b* deficiency (Figure 4C). Our results demonstrate that minor *de novo* DNA methylation may participate in establishment, but not in initiation of iXCI.

**Figure 4.**
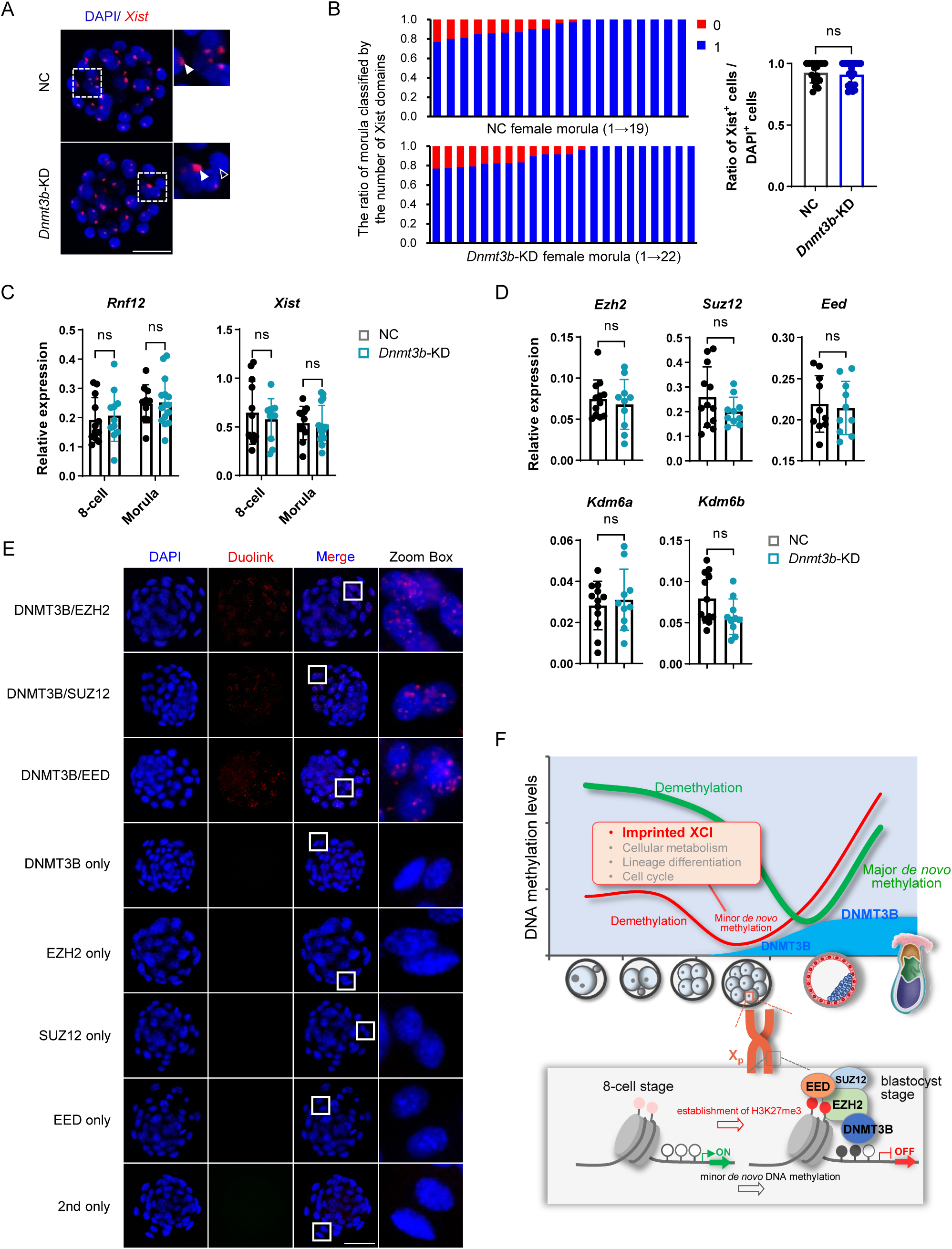
Minor *de novo* DNA methylation co-regulates iXCI via the interaction between DNMT3B and PRC2 core components. (A) Representative localization of *Xist* expression detected by RNA-FISH in the nuclei (DAPI) of NC and *Dnmt3b*-KD female morulae. (B) The ratio of blastomere with *Xist* signal to the total number of blastomeres in each NC and *Dnmt3b*-KD female morulae. Each bar represents one female embryo. The arrowhead indicates *Xist* RNA domain and the blank arrowhead indicates the blastomere without *Xist* RNA domain. (C) *Rnf12* and *Xist* expression levels in individual *Dnmt3b*-KD female 8-cell embryos and morulae. (D) Expression of H3K27me3 modifiers in individual *Dnmt3b*-KD female morulae. (E) Validation of the interaction between DNMT3B and PRC2 core components, *i.e.,* EZH2, SUZ12, EED, via *in situ* PLA. All data are presented as the mean ± SD of at least three independent experiments. \*\**P* < 0.01. ns, not significant. Scale bars: 50 µm (A, E). (F) A model illustrating a wave of minor *de novo* DNA methylation that initiates during the transition from the mouse 8-cell to blastocyst stage. Minor *de novo* DNA methylation co-regulates iXCI via the interaction between DNMT3B and PRC2 core components, and fine-tunes the processes of lineage commitment, proliferation and metabolic homeostasis before blastocyst formation.

In attempt to further understand the mechanism underlying the role of minor *de novo* DNA methylation in establishing iXCI, we next asked whether the *Dnmt3b* deficiency-induced loss of H3K27me3 domains was related to abnormal expression of histone-modifiers. We detected the expression of H3K27 histone methyltransferases or the partner (*Ezh2*, *Suz12, Eed*) and demethylase (*Kdm6a*, *Kdm6b*). The results showed *Dnmt3b* deficiency has no impact on the expression levels of these chromatin modifiers (Figure 4D).

DNA methylation and H3K27me3 have been shown to have complex relationship, possibly in a cell-type and/or genomic region-specific manner (King et al., 2016). Under certain circumstances, DNA methylation shows antagonistic effect to H3K27me3 at promoters, via excluding the binding of PRC2 components to their targets (Bartke et al., 2010; Jermann et al., 2014), while other studies have presented alternative evidence that PRC2 and DNMT cooperate to achieve silencing (Vire et al., 2006), and has a synergistic relationship in the process of global heterochromatin formation (Hagarman et al., 2013; King et al., 2016). A recent work reported that maintenance of noncanonical imprinting requires maternal allele-specific *de novo* DNA methylation in extraembryonic cells (Chen et al., 2019). It has been thought that during the late phase of rXCI in fully differentiated cells, gene silencing is initially achieved by PRC2 complex-induced H3K27me3, and further stably maintained by the redundant action of multiple layers of epigenetic modifications, including DNA methylation, to reach the maximum level of chromatin compaction (Chow and Heard, 2009; Heard et al., 2004; Pintacuda and Cerase, 2015). In line with this, a recent multifaceted analysis showed that DNA methylation and H3K27me3 are concurrently enriched in genes subject to XCI (Balaton and Brown, 2021). Thus, we next reanalyzed H3K27me3 dynamics of promoters that underwent minor *de novo* DNA methylation during the transition from the 8-cell to ICM, using a well-established H3K27me3 ChIP-seq data (Liu et al., 2016). As expected, concurrent increase in H3K27me3 enrichment can be clearly observed throughout the X chromosome and many *de novo* methylated X-linked promoters (Figure 4—figure supplement 2A, B). Next, we used *in situ* proximity ligation assays (PLA), a method that is suitable for visualizing protein interactions by detecting the close proximity of two molecules (Soderberg et al., 2006) to test interactions between DNMT3B and PRC2 core components, *i.e.,* EZH2, SUZ12, EED, which are essential for X-linked heterochromatin. It is clearly visible that DNMT3B-EZH2, DNMT3B-SUZ12 and DNMT3B-EED interactions can be detected in the nuclei by the blastocyst stage (Figure 4E). In line with this, the function of *de novo* DNA methylation in XCI, as well as direct interaction between DNMT3B and PRC2 core components, were also confirmed in differentiated female ES cells (Figure 4—figure supplement 3A-H). Collectively, our results support the idea that minor *de novo* DNA methylation plays an important role in iXCI via synergistic interaction with heterochromatic histone modifications, rather than transcriptional regulation of iXCI-regulating genes.

It is also worth mentioning that minor *de novo* DNA methylation is definitely distinct from previously reported DNA re-methylation in monkey and human preimplantation embryos, which occurs during the 2- to 8-cell and 4- to 8-cell transition respectively (Gao et al., 2017; Guo et al., 2014; Zhu et al., 2018). The so-called DNA re-methylation does not occur in naturally conceived mouse embryos. However, an aberrant wave of DNA re-methylation has been identified in cloned mouse preimplantation embryos as an epigenetic barrier that impairs full-term developmental potential (Gao et al., 2018).

Collectively, our study presents an update on the current knowledge of embryonic *de novo* DNA methylation. We identified that a wave of minor *de novo* DNA methylation has initially occurred during the transition from the mouse 8-cell to blastocyst stage, but not during the implantation stage. Of interest, we provide the evidence that DNA methylation is important for establishing iXCI in preimplantation female embryos. Our data, together with previous results, also support a model where minor *de novo* DNA methylation and demethylation co-orchestrate DNA methylation homeostasis before blastocyst formation, and serve as epigenetic mechanisms affecting embryonic developmental potential by fine-tuning the processes of lineage commitment, proliferation and metabolic homeostasis (Figure 4F). Thus, our work provides a novel insight for understanding epigenetic programming and reprogramming during early embryonic development.

## MATERIALS AND METHODS

### Mouse embryo preparation

All experimental procedures were approved by and performed in accordance with the guidelines of the Institutional Animal Care and Use Committee of China Agricultural University. Mice were fed ad libitum and housed under controlled lighting (12 light:12 dark) and specific pathogen-free conditions. ICR female mice were superovulated with 5 IU pregnant mare serum gonadotrophin (PMSG; Ningbo Hormone Product Co., Ltd, Ningbo, China) and a further 5 IU human chorionic gonadotropin (hCG; Ningbo Hormone Product CO., Ltd) 46-48 h later. Then, superovulated ICR females were cogaged individually with ICR males. *In vivo* zygote, 2-cell embryos, 4-cell embryos, 8-cell embryos, morulae, or blastocysts were flushed from oviducts and uterus at 16 h, 42 h, 54 h, 66 h, 78 h, or 93 h post-hCG, respectively.

The *Dnmt3b*^tm2Enl^/^+^ mouse line(Okano et al., 1999) was purchased from Mutant Mouse Resource & Research Centers (MMRRC, 029886-UNC). For collection of *Dnmt3b*^+^/ ^+^ (WT) and *Dnmt3b* ^tm2Enl^/ ^tm2Enl^ (KO) embryos, *Dnmt3b*^tm2Enl^/^+^ female mice were superovulated as above, and mated with *Dnmt3b*^tm2Enl^/^+^ male mice. Morulae and blastocysts were collected at 78 h and 93 h, respectively.

### Quantitative real-time PCR analysis

Total RNA was extracted from embryos using TRIzol^TM^ (Invitrogen, Carlsbad, CA, USA). RNA was treated with DNase, and reverse-transcribed into cDNA using the HiScript II Q Select RT Kit (Vazyme, Nanjing, China). qRT-PCR was performed using SsoFast Eva Green Supermix (Bio-Rad, Hercules, CA, USA). The relative expression data were calculated using the 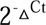 method. Relative expression level was normalized to the housekeeping gene *H2afz*, and each experiment was repeated at least three times, as previously reported. Primers used are listed in Table S1.

For qRT-PCR on single female or male embryo, a blastomere was isolated and treated with 2 μL Cells-to-cDNA™ II Cell Lysis Buffer (Invitrogen) at 75°C for 15 min. Next, the lysate was digested with proteinase K (Merck, Kenilworth, NJ, USA) at 55°C for 2 h, and used for sex determination according to the manufacturer’s instruction of Ex Taq Hot Start Version (Takara Bio, Shiga, Japan). The amplification was conducted as follows: 95 °C for 3 min; 40 cycles of 95 °C for 10 s, 52 °C for 15 s, 72 °C for 15 s; and extension at 72 °C for 5 min. The remaining part of embryo was treated with 5 μL Cells-to-cDNA™ II Cell Lysis Buffer (Invitrogen), and then reverse-transcribed into cDNA with the HiScript II Q Select RT Kit (Vazyme).

### Immunofluorescence analysis

Embryos were fixed with 4% Paraformaldehyde (PFA) in PBS-0.1% PVA at 4 °C overnight. After permeabilizing with 0.5% Triton X-100 in PBS-0.1%PVA (PBST-PVA) for 1 h at room temperature, embryos were blocked with 1% BSA in PBST-PVA at 4 °C for 6 h. Embryos were incubated with primary anti-DNMT3B antibody (1:200; GeneTex, Irvine, CA, USA), anti-H3K27me3 antibody (1:1000; MilliporeSigma, Burlington, MA, USA), or anti CDX2 antibody (1:200; BioGenex, Fremont, CA, USA) overnight at 4 °C, followed by Alexa Fluor-488 (anti-mouse; Invitrogen) and Alexa Fluor-594 (anti-rabbit; Invitrogen) labeled secondary antibodies for 1 h at room temperature. Finally, embryos were counterstained with DAPI. Fluorescence images were obtained under an upright microscope (BX51; Olympus) using an attached digital microscope camera (DP72; Olympus). Quantification of the immunofluorescence results were performed using Image J (National Institutes of Health, Bethesda, MD, USA).

For 5-methylcytosine (5-mC) staining, permeabilized embryos were treated with 4 M HCl for 10 min and 100 mM Tris-HCl for 10 min at room temperature, and then blocked overnight at 4 °C. Next, embryos were incubated with primary anti-5mC antibody (1:200; Active Motif, Carlsbad, CA, USA) at room temperature for 2 h, followed by secondary antibody for 1 h.

Furthermore, embryos for sex determination and genotyping were removed from the slides and digested with Cells-to-cDNA™ II Cell Lysis Buffer (Invitrogen) and proteinase K (Merck) as described above, followed by sex determination and genotyping using PCR. For genotyping, the amplification was conducted as follows: 94 °C for 3 min; 40 cycles of 94 °C for 30 s, 65 °C for 30 s, 72 °C for 45 s; and extension at 72 °C for 7 min. The primer sequences are listed in Table S1.

### Bisulfite sequencing

The DNA methylation profile was analyzed by PCR-based bisulfite sequencing. 40 IVO blastocysts or 80 8-cell embryos were digested and bisulfite converted according to the manufacturer’s instruction of EZ DNA Methylation-Direct Kit (Zymo, Irvine, CA, USA). Bisulfite-specific primers were designed with the online Methyl Primer Express, v1.0 (http://www.urogene.org/methprimer/ Table S1). The converted DNA was amplified by PCR using Ex Taq Hot Start Version (Takara). And the PCR products were subcloned into pEASY-T5 Zero Cloning Kit (TransGen, Beijing, China). At least 10 clones per group were sequenced. The sequencing results were analyzed with QUMA (http://quma.cdb.riken.jp/).

### Microinjection of siRNA

siRNA oligo sequences were synthesized by GenePharma (Shanghai, China). The sequences of *Dnmt3b* siRNA used in the present study were as follows: sense 5’-CCUCAAGACAAAUAGCUAUTT-3’, antisense 5’-AUAGCUAUUUGUCUUGAGGTT-3’. siRNA was diluted to 100 μM and stored at -80 °C. For siRNA microinjection, 5 pL siRNA was microinjected into the cytoplasm of zygotes. Then embryos were cultured in KSOM at 37 °C under 5% CO_2_ and collected at the 8-cell, morula or blastocyst stage for further analyses.

### Treatment of 5-aza-dC

Zygotes were collected as described above, and 10 nM 5-aza-2’-deoxycytidine (5-aza-dC) was supplemented into *in vitro* culture medium from the 8-cell stage onward. Blastocysts were collected at 110 h post-hCG administration.

### RNA-FISH

RNA-FISH was performed using ViewRNA™ ISH Cell Assay Kit (Invitrogen) according to the manufacturer’s instructions. Fixed embryos were treated with Detergent Solution QC at room temperature for 5 min, and digested with protease QS at room temperature for 10 min. Next, the embryos were hybridized with probes against *Xist* (Invitrogen) at 40 °C for 3 h. Then the embryos were hybridized with PreAmplifier and Amplifier Mix at 40 °C for 30 min, respectively. Label Probe Mix was used to produce a signal. The embryos were counterstained with DAPI, and imaged using an attached digital microscope camera (DP72; Olympus).

### Proximity ligation assays

The in situ proximity ligation assays (PLA) was performed using the Duolink® In Situ Red Starter Kit according to the manufacturer’s instruction (MilliporeSigma). Embryos were fixed and permeabilized as described for immunofluorescence. After blocking, embryos were incubated with primary antibody. Embryos were then incubated with PLA probes, followed by ligation and amplification reaction. Finally, embryos were counterstained with Duolink® In Situ Mounting Medium with DAPI (MilliporeSigma). Furthermore, DNMT3B, EZH2, SUZ12, or EED single primary antibody, and secondary antibodies alone were used as technical negative controls.

### Generation of Dnmt3a/Dnmt3b KO ESC

Mouse ESCs line PGK12.1 was used in the present study. Cells were cultured in KnockOut™ DMEM medium (Invitrogen) containing 10% fetal bovine serum (VISTECH, Sydney, Australia), 2mM GlutaMAX (Invitrogen), 1×non-essential amino acid (MilliporeSigma), 55 μM β-mercaptoethanol (Invitrogen), 1000 units/ml mouse recombinant leukemia inhibitor factor (MilliporeSigma), 100 units/ml penicillin and 100 µg/ml streptomycin (both Invitrogen). To induce ESC differentiation, ESCs were cultured in DMEM/F12 (Invitrogen) and Neurobasal medium supplement with B27 and N2 supplement (Invitrogen), 100 units/ml penicillin-streptomycin (Invitrogen).

Knockout of *Dnmt3b* or *Dnmt3a* for ES cells was performed using CRISPR/Cas9n. sgRNA was designed using Benchling (https://www.benchling.com/). sgRNAs were constructed to pSpCas9n(BB)-2A-Puro vector (PX462, Addgene plasmid no. 62987). 4 μg of the plasmid was transfected using Lipofectamine 2000 (Invitrogen) according to the manufacturer’s protocol. 2 μg/ml puromycin (Solarbio, Beijing, China) was used to selected the positive colonies. After selection, colonies were picked out and expanded. Cells were verified by genotyping and Sanger sequencing. The sgRNA sequences and knockdown cell detection primers are listed in Table S1.

### Co-immunoprecipitation

Co-immunoprecipitation (Co-IP) was performed with Dynabeads™ Protein G Immunoprecipitation Kit (Invitrogen) according to the manufacturer’s protocol. Briefly, 3 μg DNMT3B antibody (GeneTex) or mouse IgG (MilliporeSigma) was incubated to Protein G beads with rotation for 10 minutes at room temperature. The beads were washed with Washing Buffer. Then cell lysates were then incubated with the beads-antibody complex with rotation for 4h at 4 °C. Then the beads were washed 3 times with Washing Buffer, and were eluted with loading buffer by heating at 98 °C for 10min. Co-IP products were analyzed by western blot using the indicated primary antibodies and secondary antibodies. And protein bands were detected with Tanon 5200 detection systems (Tanon, Shanghai, China).

### Bioinformatics analysis

RRBS data of mouse embryos (2-cell embryos, 4-cell embryos, 8-cell embryos, ICM, and E6.5 embryos) were downloaded from the published article by Smith et al (Smith et al., 2012b) (accession number: GSE34864). The methylation level was calculated as the number of “methylated” reads (reporting as C), divided by the total number of “methylated” and “unmethylated” read, which reporting as C or T. The genomic region information was downloaded from the mm9 Repeat Masker. As described in the published article, promoters were defined as 1 kb up- and downstream of the TSS and classified into high-density CpG promoter (HCP), intermediate-density CpG promoter (ICP) and low-density CpG promoter (LCP). Only CpG sites with at least fivefold coverage were included in the methylation analysis. WGBS data of 8-cell and ICM were from GSE84236. Cool-seq data of female and male embryos were from GSE78140, and analyzed as described in the article (Guo et al., 2017). Briefly, every covered WCG site (W includes A or T) with at least one-fold coverage was summed for the analysis of DNA methylation levels. The bigwig files of RRBS, RNA-seq and H3K27me3 CUT&RUN data of WT and Dnmt3a/3b DKO E6.5 ExE were downloaded from the published article by Chen et al (Chen et al., 2019) (accession number: GSE130115). The bigwig files of H3K27me3 ChIP-seq data of 8-cell embryos and ICM were downloaded from the published article by Liu et al. (Liu et al., 2016) (accession number: GSE73952).

Gene ontology (GO) annotation analysis was performed using Database for Annotation, Visualization and Integrated Discovery (DAVID 6.8: https://david.ncifcrf.gov/). Heatmaps were plotted using the R software “pheatmap” package (version 1.0.12). The integrative genomics viewer (IGV) was used to visualize the ChIP-seq results. The phenotype of genes was analyzed on the Mouse Genome Informatics (MGI; http://www.informatics.jax.org/) database.

### EdU assay

The proliferation of embryos was detected with a BeyoClick™ EdU-594 Cell Proliferation Kit (Beyotime, Shanghai, China) according to the manufacturer’s instruction. Embryos were incubated in EmbryoMax® KSOM Mouse Embryo Media (MilliporeSigma) containing 10 μM EdU at 37 °C for 2 h, and then fixed and permeabilized. Afterwards, embryos were incubated with the Click Reaction mixture for 2 min at room temperature before stained with DAPI. Images were acquired under an upright microscope (BX51; Olympus) using an attached digital microscope camera (DP72; Olympus).

### ATP measurement

ATP level was measured using an ATP determination kit according to the manufacturer’s instruction (Beyotime). Briefly, blastocysts were incubated with M2 medium, and 1 μM oligomycin (Abmore, Houston, USA) or 1 mM sodium oxamate (MilliporeSigma) in M2 medium at 37 °C for 15 min, respectively. Next, embryos were digested with 20 μL ATP lysis buffer on ice, and the lysate was added to 96-well plates containing 100 μL ATP detection working dilution. The luminescence was detected by a multifunctional microplate reader (Infinite F200; TECAN, Mannedorf, Switzerland).

### Embryo transfer

Pseudo-pregnant ICR females were mated with ICR males 3.5 d before embryo transfer. Blastocysts derives from NC or *Dnmt3b*-KD group were transferred to the uterine horn of pseudo-pregnant female mice.

### Statistical analysis

All data are presented as mean ± SD and analyzed with the student’s *t* test or correct χ^2^ procedure by using SPSS v.25.0 (IBM, Armonk, NY, USA). Values of *P* < 0.05 were considered statistically significant.

## Data availability

All data are available in the article and/or supporting information.

## Acknowledgements

We thank Neil Brockdorff (University of Oxford) and Ingolf Bach (University of Massachusetts Medical School) for the gift and delivery of PGK12.1 cells. We thank Zihuan Du for the assistance in the analysis of sequencing data. This work was supported by grants from the National Natural Science Foundation of China (31930103), National Agriculture Key Science & Technology Project (NK20221201), Ningbo Major Science and Technology Project (2021Z112), the National Key R&D Program (2022YFD1300301).

## Author contributions

Y.Y., J.T. and L.A. conceived and designed the experiments. Y.Y. C.Z. and W.W. were responsible for animal care and management. Y.Y., W.F., Q.Y., C.Z., W.W., M.C., Q.L., Y.T., J.C., X.W. and Z.Z. performed the experiments. Y.Y., W.F., J.T. and L.A. analyzed the data. Y.Y., W.F., J.T. and L.A wrote, and finalized the manuscript. All authors approved the paper.

## Supplemental Tables (separate files)

**Supplemental Table S1.** Primers sequences used in this study.

## Figure legends

**Figure 1—figure supplement 1.**
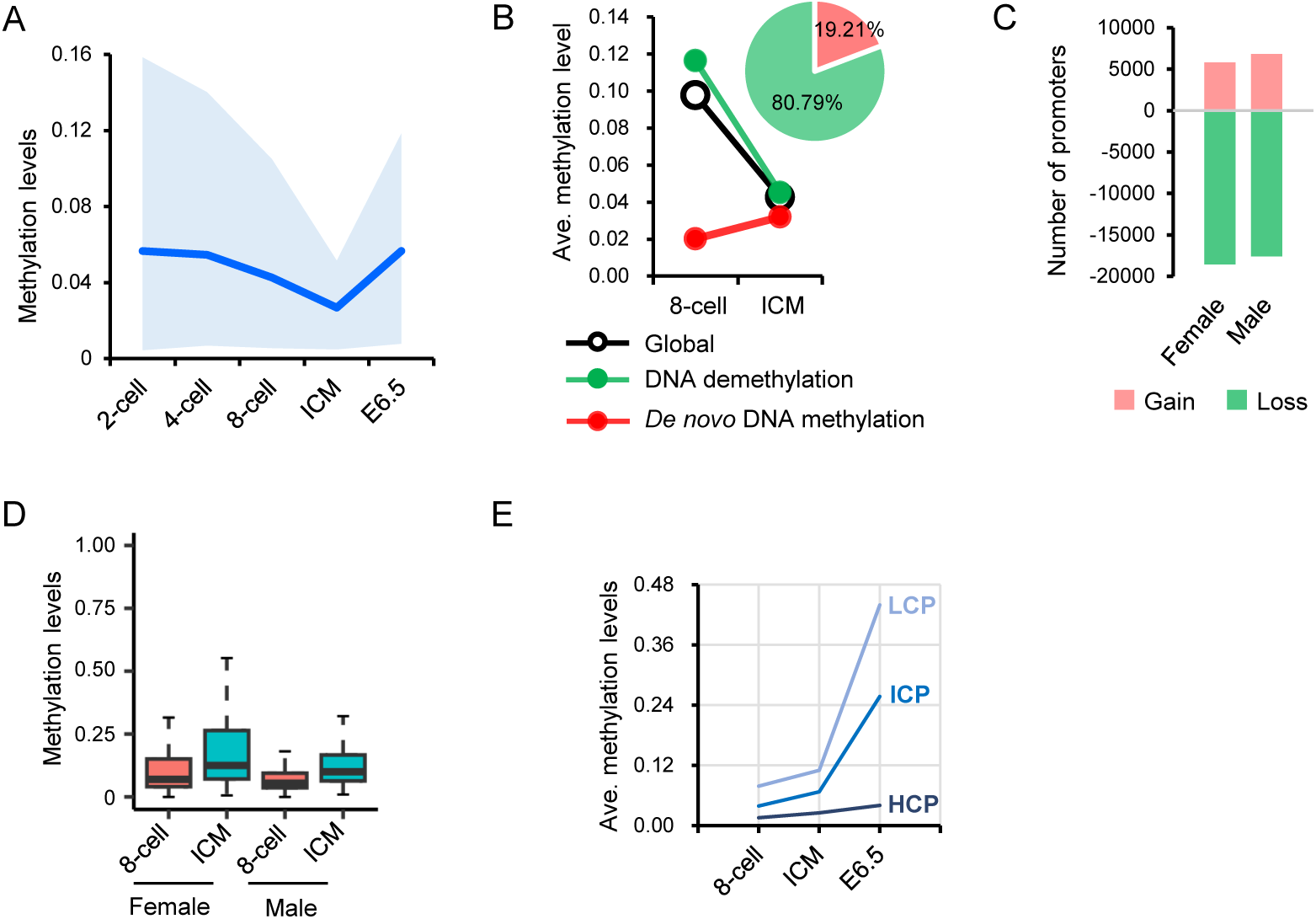
(A) Dynamics of global methylation during preimplantation embryos (blue line, average methylation levels; colored space, 10th/ 90th percentile). (B) General trends of CpG islands undergoing *de novo* methylation or demethylation, and their relative contribution to differentially methylated regions. (C) Number of promoters that gain or loss DNA methylation in female and male embryos during the transition from the 8-cell to blastocyst stage. (D) DNA methylation changes of promoters that gain DNA methylation during the transition from the 8-cell to blastocyst stage in female or male embryos. (E) DNA methylation dynamics of promoters with different CpG-densities during consecutive transitions.

**Figure 2—figure supplement 1.**
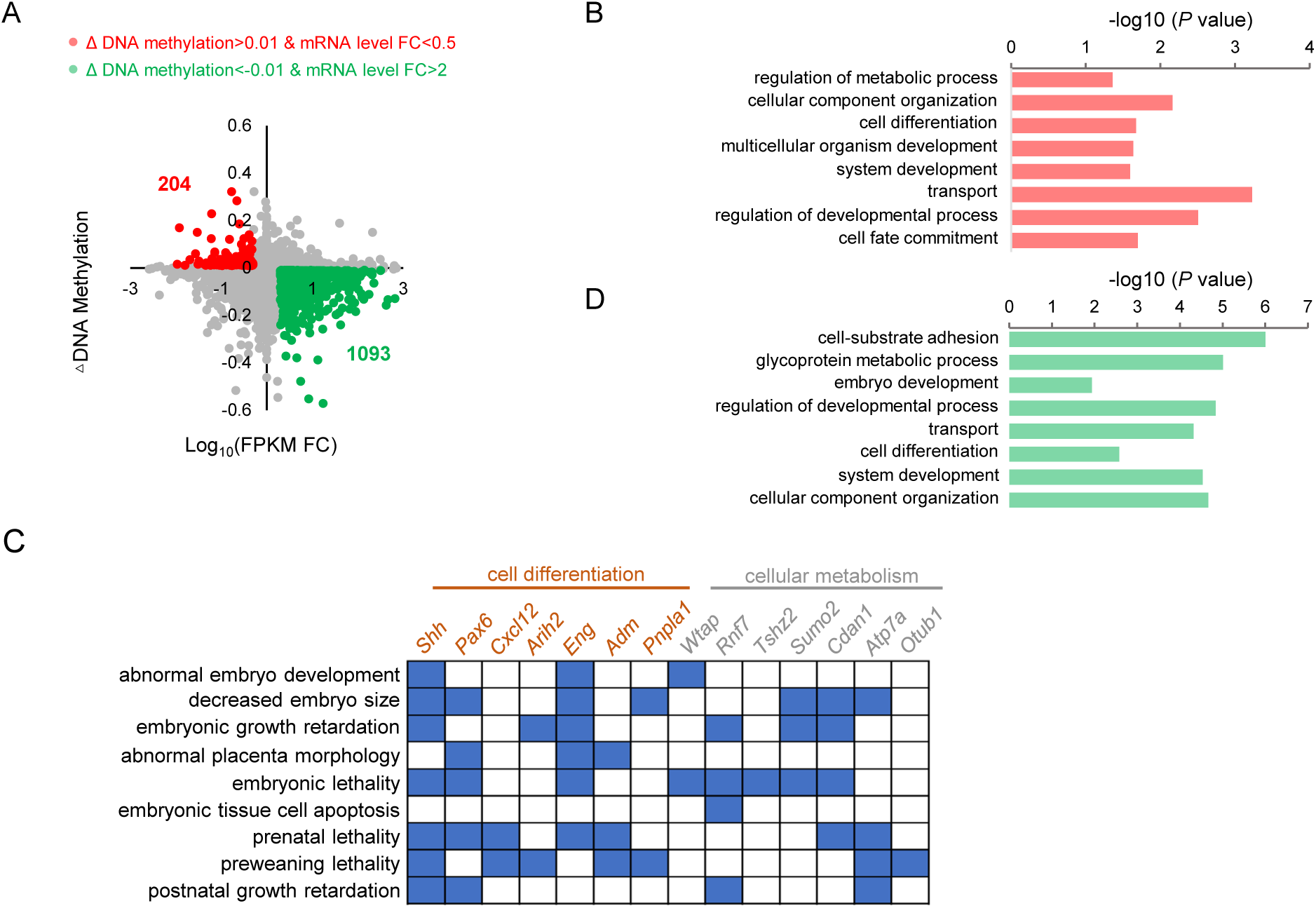
(A) A scatter plots of changes in promoter DNA methylation and gene expression from the 8-cell embryos to ICM. Red plots indicate putative genes that undergo minor *de novo* DNA methylation (ΔDNA methylation > 0.01) and are downregulated (ΔDNA methylation < -0.01). (B-C) Gene ontology analysis (B) and phenotype annotation (C) of genes transcriptionally regulated by minor *de novo* DNA methylation. Blue squares indicate genes that are related to developmental or lethal phenotypes. (D) Gene ontology analysis of genes transcriptionally regulated by DNA demethylation from the 8-cell embryos to ICM.

**Figure 2—figure supplement 2.**
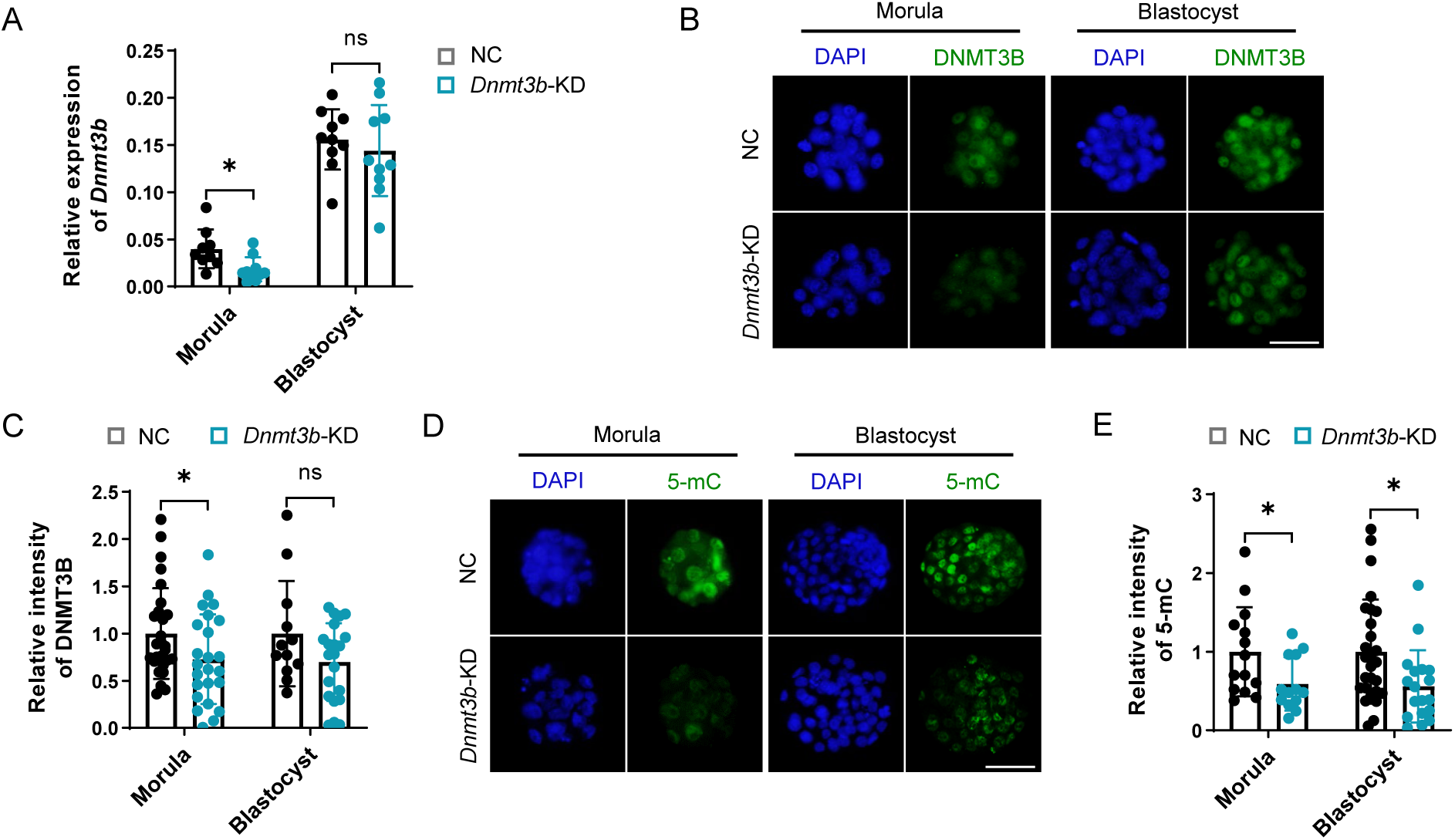
(A) Relative expression levels of *Dnmt3b* in NC and *Dnmt3b*-KD morulae and blastocysts. (B-C) Immunofluorescent staining (C) and quantification (D) of DNMT3B in NC and *Dnmt3b*-KD morulae and blastocysts. (D-E) Immunofluorescent staining (D) and quantification (E) of 5-mC in NC and *Dnmt3b*-KD morulae and blastocysts. All data are presented as the mean ± SD of at least three independent experiments. \**P* < 0.05, \*\**P* < 0.01. ns, not significant. Scale bars: 50 µm (B, D).

**Figure 2—figure supplement 3.**
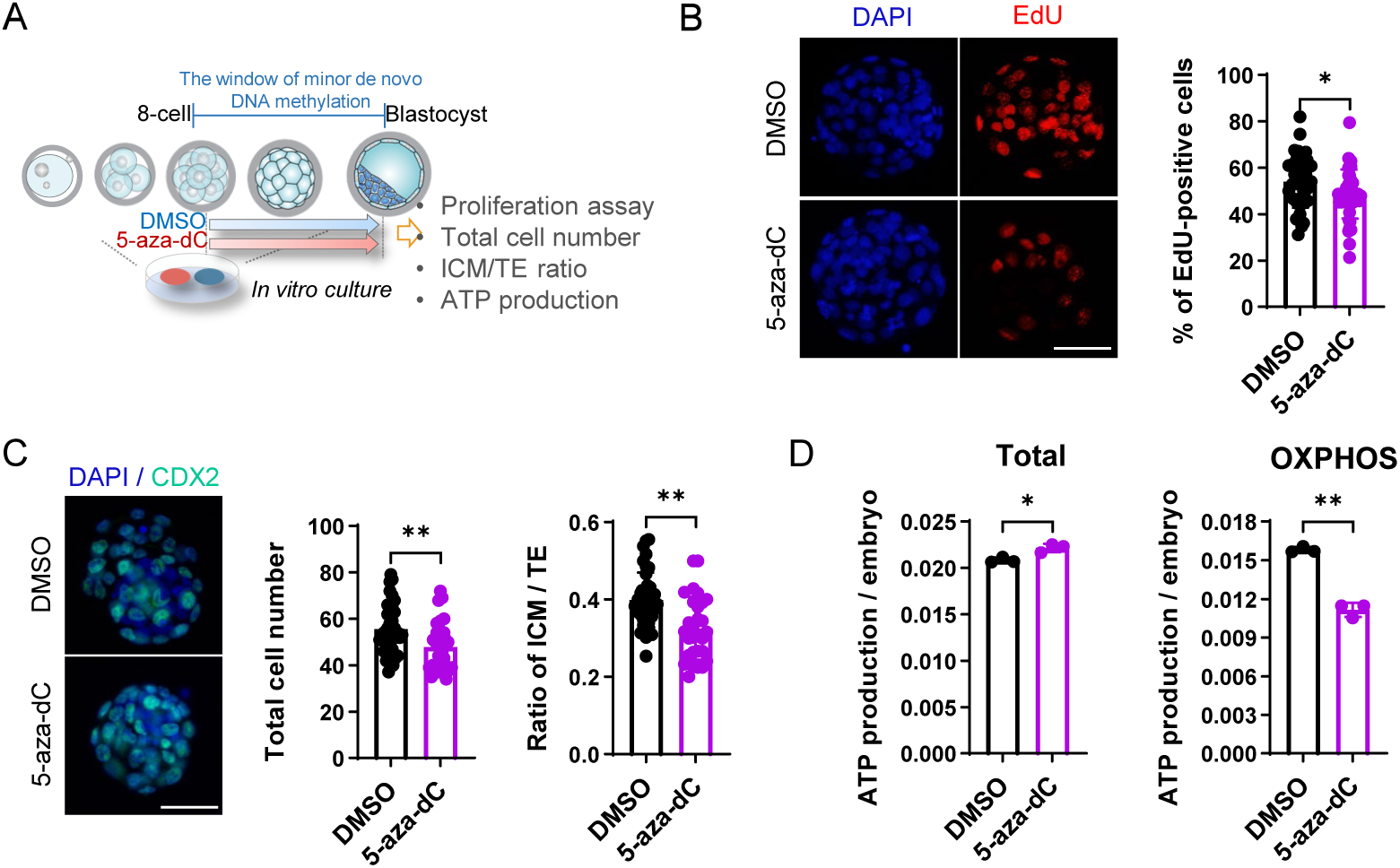
(A) Schematic diagram of the experimental design. (B) EdU staining of blastocysts exposed to 5-aza-dC from the 8-cell to blastocyst stage. Right panel: the percentage of EdU-positive cells. (C) The total cell number (left panel) and the ratio of ICM/TE (right panel) of blastocysts treated with or without 5-aza-dC. (D) The total ATP production (left panel) and OXPHOS-dependent ATP production (right panel) of blastocysts treated with or without 5-aza-dC. All data are presented as the mean ± SD of at least three independent experiments. \**P* < 0.05, \*\**P* < 0.01. ns, not significant. Scale bars: 50 µm (B, C).

**Figure 3—figure supplement 1.**
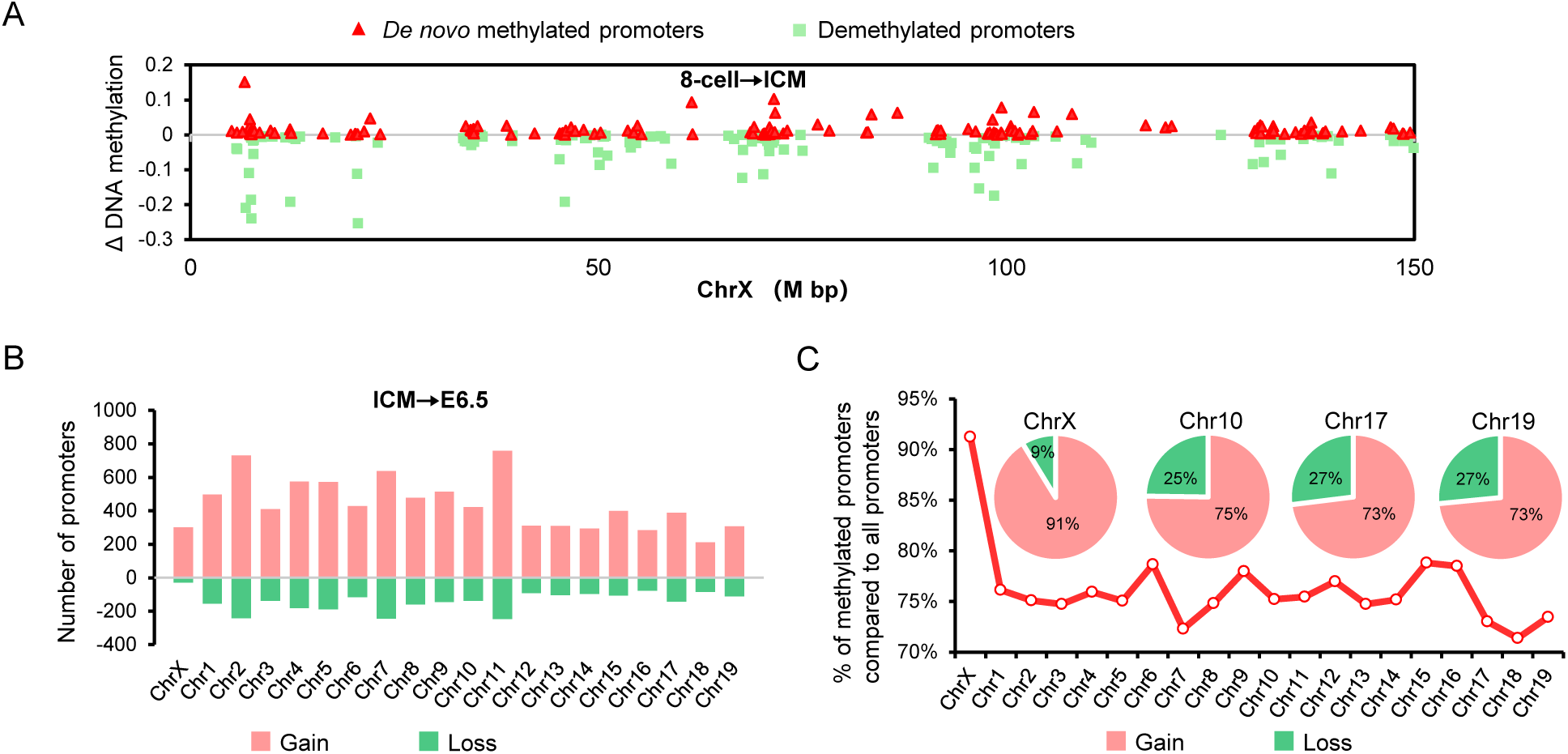
(A) The distribution of *de novo* methylated and demethylated promoters cross the X chromosome during the transition from the 8-cell to blastocyst stage. (B) The chromosome-wide distribution of *de novo* methylated promoters during the transition from the blastocyst to post-implantation stage. (C) Percentage of *de novo* methylated promoters on each chromosome during the transition from the blastocyst to post-implantation stage.

**Figure 3—figure supplement 2.**
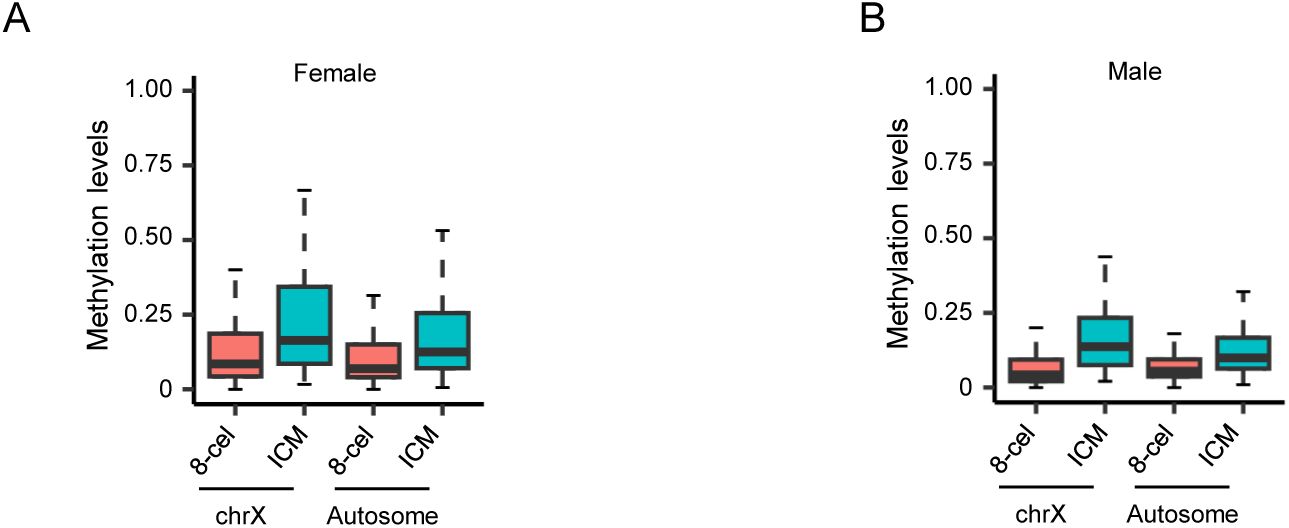
(A) DNA methylation changes of promoters that gain DNA methylation during the transition from the 8-cell to blastocyst stage in chromosome X and autosomes of female embryos. (B) DNA methylation changes of promoters that gain DNA methylation during the transition from the 8-cell to blastocyst stage in chromosome X and autosome of male embryos.

**Figure 3—figure supplement 3.**
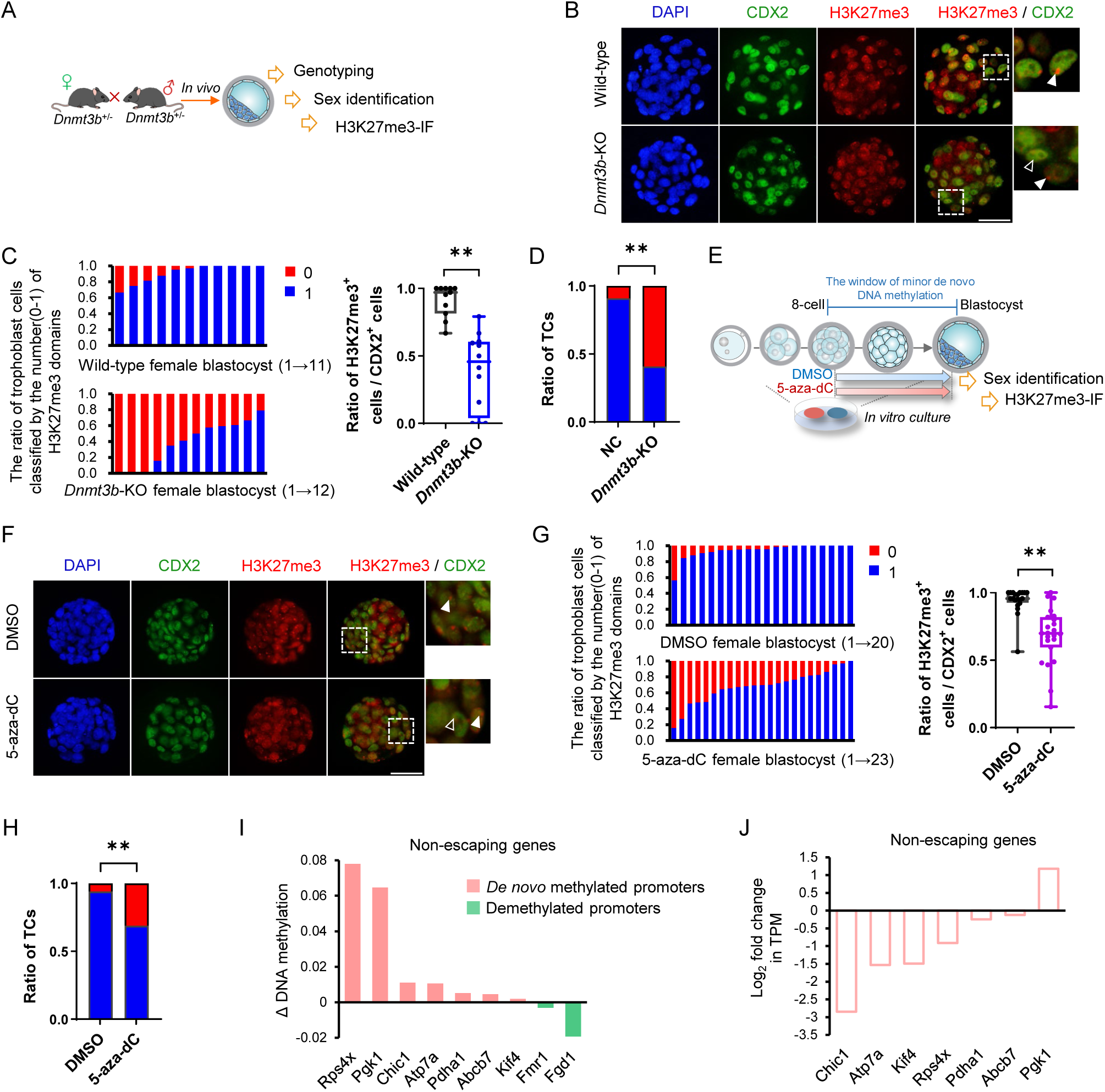
(A) Schematic diagram of the experimental design. *Dnmt3b*-KO female embryos were collected by intercrossing *Dnmt3b*^+/-^ male and female mice. Immunostaining assay for H3K27me3 was performed and quantified in trophoblast cells in each female homozygous blastocyst. (B) Representative immunostaining for H3K27me3 (red) in the nuclei (DAPI) of wild-type and *Dnmt3b*-KO female blastocysts colabeled with CDX2 (green)-positive trophoblast cells. The arrowhead indicates the H3k27me3 domain and the blank arrowhead indicates the blastomere without the H3k27me3 domain. (C) The ratio of trophoblast cells classified by the number of H3K27me3 domains in each wild-type and *Dnmt3b*-KO female blastocysts. Each bar represents one female embryo. (D) The percentage of H3K27me3-positive and H3K27me3-negative trophoblast cells (TCs) to total trophoblast cells in female blastocysts. (E) Schematic diagram of the experimental design. Zygotes were cultured in vitro and transiently exposed to 5-aza-dC during the window of minor *de novo* DNA methylation (the 8-cell to blastocyst stage) or not. Immunostaining assay for H3K27me3 was performed and quantified in trophoblast cells in each female blastocyst. (F) Representative immunostaining for H3K27me3 (red) in the nuclei (DAPI) of 5-aza-dC treated or untreated female blastocysts colabeled with CDX2 (green)-positive trophoblast cells. Scale bars: 50 µm. The arrowhead indicates the H3k27me3 domain and the blank arrowhead indicates the blastomere without the H3k27me3 domain. (G) The ratio of trophoblast cells classified by the number of H3K27me3 domains in female blastocysts exposed or not exposed to 5-aza-dC from the 8-cell to blastocyst stage. Each bar represents one female embryo. (H) The percentage of H3K27me3-positive and H3K27me3-negative trophoblast cells (TCs) to total trophoblast cells in female blastocysts. (I) The changes of promoter methylation levels of X-linked non-escaping genes from the 8-cell to blastocyst stage. (J) The expression level changes of *de novo* methylated X-linked non-escaping genes during the transition from 8-cell to blastocyst stage. All data are presented as the mean ± SD of at least three independent experiments. \**P* < 0.05, \*\**P* < 0.01. ns, not significant. Scale bars: 50 µm (B, E).

**Figure 3—figure supplement 4.**
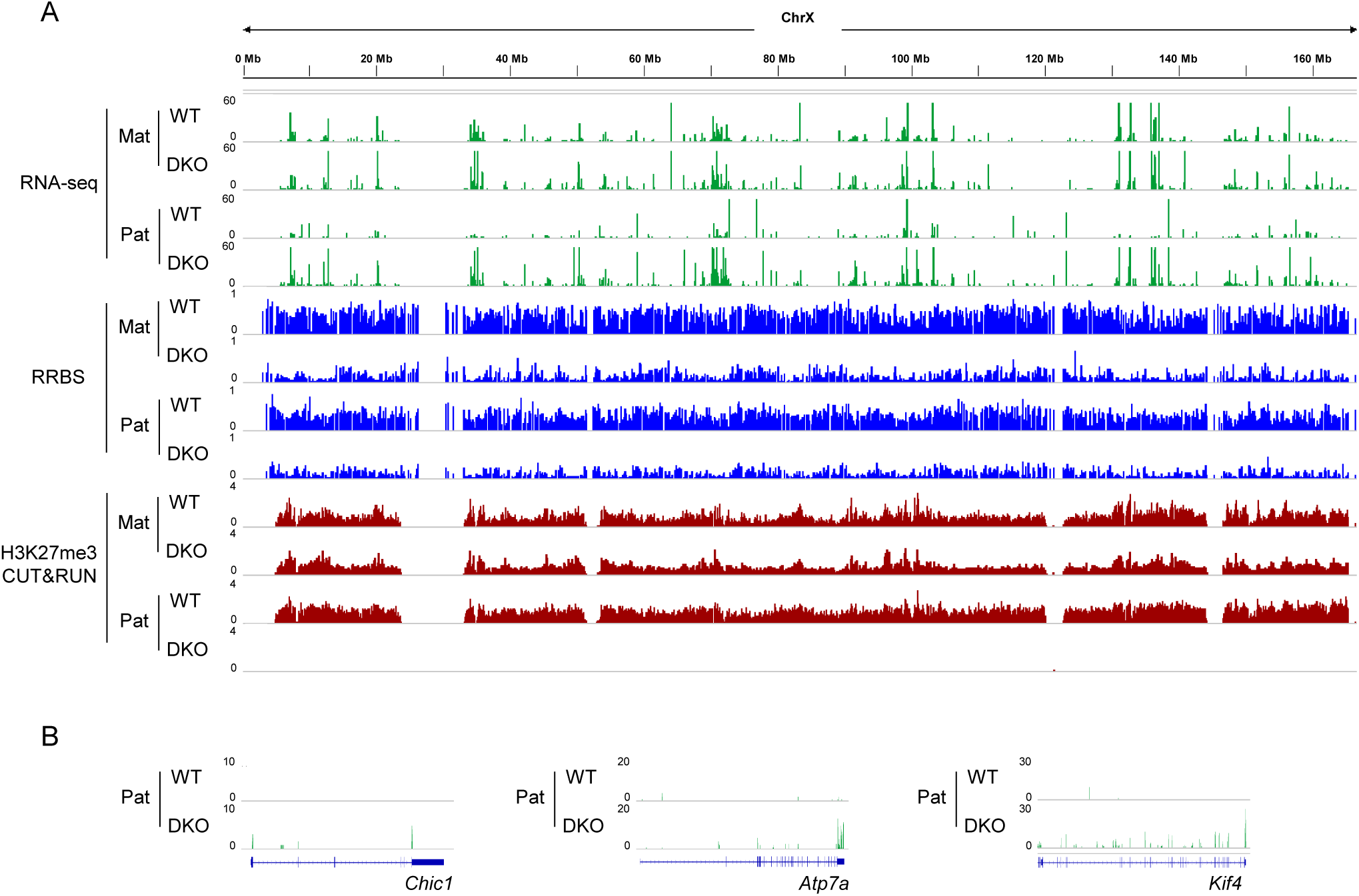
(A) Genome browser views of the allelic expression, DNA methylation and H3K27me3 enrichment cross the X chromosome in WT and DKO female ExE based on the previously published data (Chen et al., 2019). (B) Genome browser views of the paternal expression of selected X-linked non-escaping genes that undergo minor *de novo* DNA methylation based on the previously published data (Chen et al., 2019).

**Figure 4—figure supplement 1.**
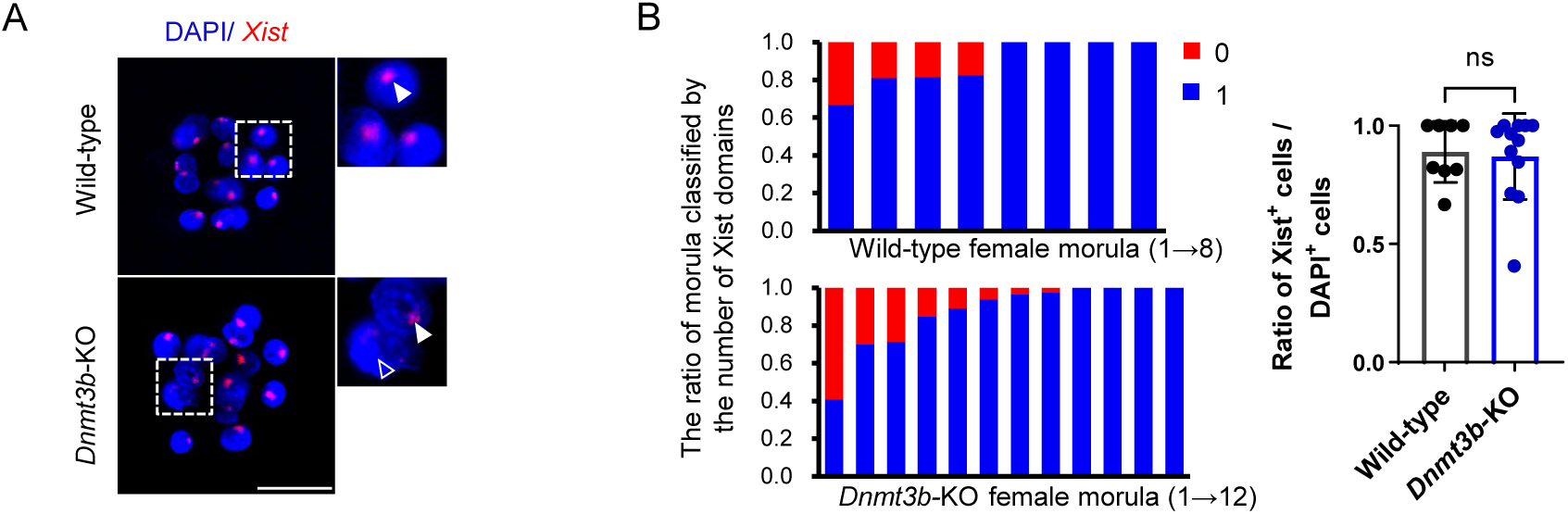
(A) Representative localization of *Xist* expression detected by RNA-FISH in the nuclei (DAPI) of wild-type and *Dnmt3b*-KO female morulae. The arrowhead indicates *Xist* RNA domain and the blank arrowhead indicates the blastomere without *Xist* RNA domain. Scale bars: 50 µm. (B) The ratio of blastomere with Xist signal to the total number of blastomeres in each wild-type and *Dnmt3b*-KO female morulae. Each bar represents one female embryo. All data are presented as the mean ± SD of at least three independent experiments. ns, not significant.

**Figure 4—figure supplement 2.**
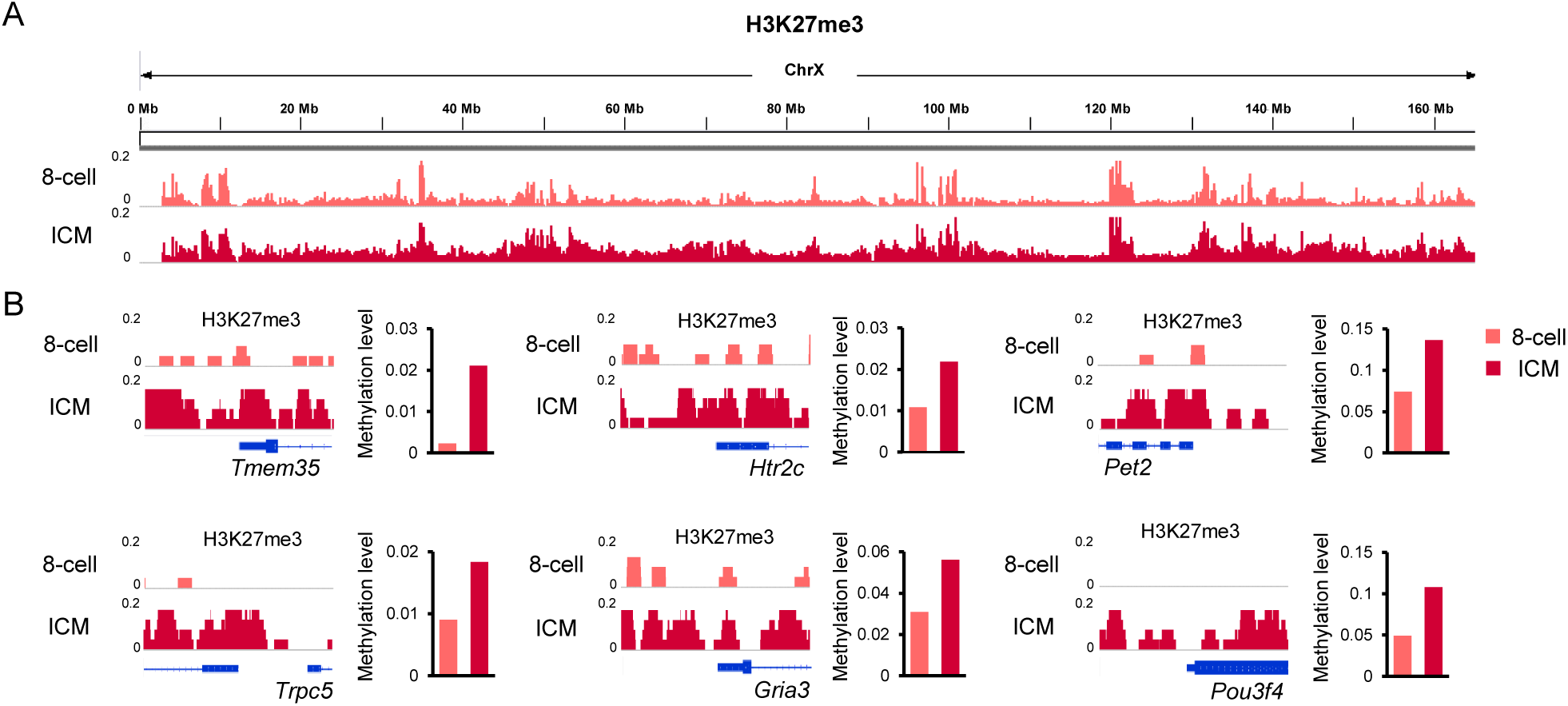
(A) H3K27me3 enrichment cross the X chromosome from the 8-cell to blastocyst stage based on the previously published data (Liu et al., 2016). (B) Concurrent increase in H3K27me3 enrichment (left panel) and DNA methylation levels (right panel) at selected X-linked promoters that undergo minor *de novo* DNA methylation.

**Figure 4—figure supplement 3.**
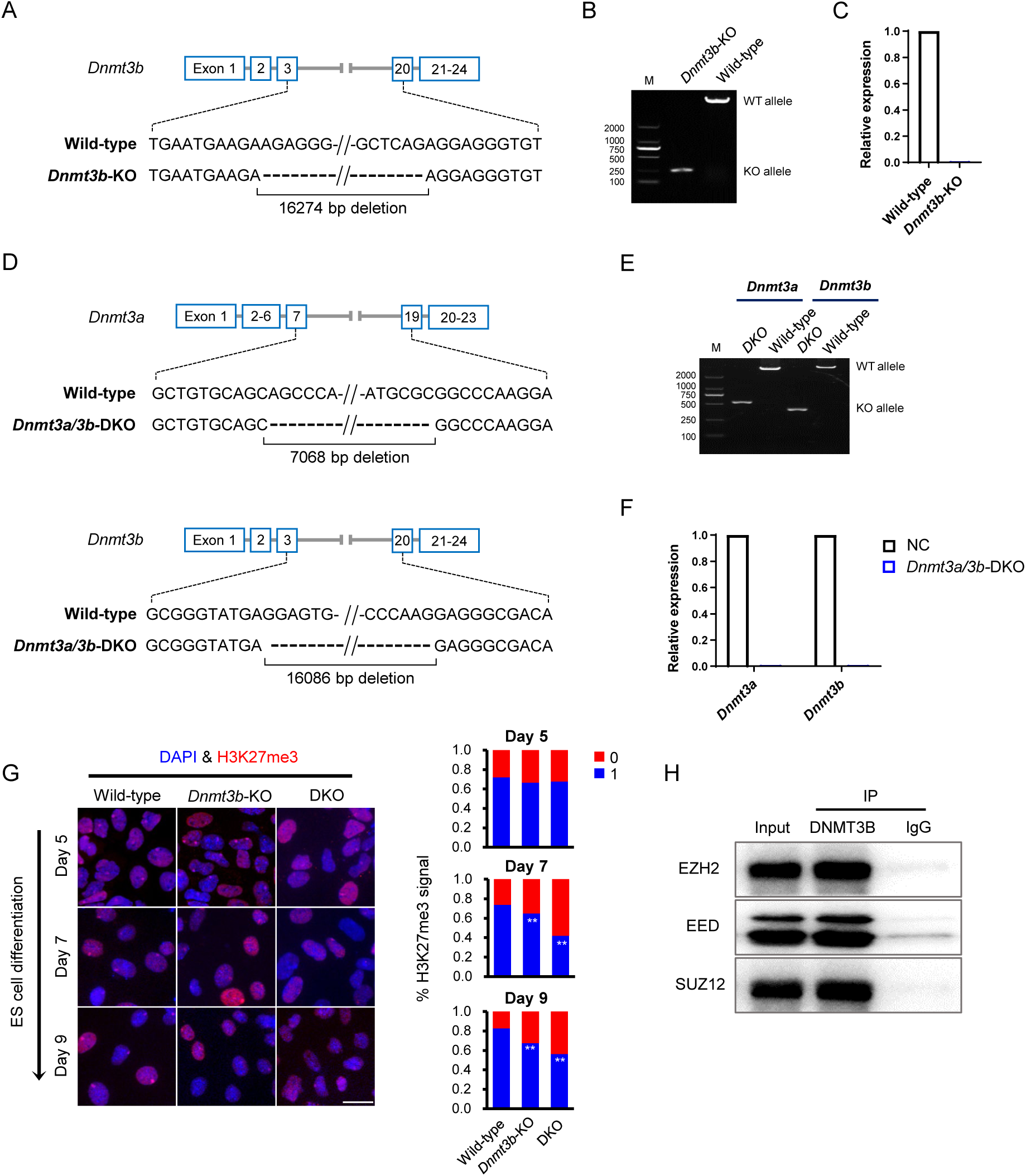
(A) Schematic illustration of generation of *Dnmt3b*-KO female ES cells via the CRISPR/Cas9n system. (B) Detection of deletion introduced by sgRNA-Cas9n targeting *Dnmt3b* via PCR with genomic DNA from wild-type and *Dnmt3b*-KO female ES cells. (C) Relative expression levels of *Dnmt3b* in wild-type and *Dnmt3b*-KO female ES cells. (D) Schematic illustration of generation of DKO female ES cells via the CRISPR/Cas9n system. (E) Detection of deletion introduced by sgRNA-Cas9n targeting *Dnmt3a/3b* via PCR with genomic DNA from wild-type and DKO female ES cells. (F) Relative expression levels of *Dnmt3a* and *Dnmt3b* in wild-type and DKO female ES cells. (G) Representative immunostaining for H3K27me3 (red) in the nuclei (DAPI) of wild-type, *Dnmt3b*-KO, and DKO female ES cells differentiated for 5, 7 or 9 days, as well as the ratio of ES cells classified by the number of H3K27me3 domains. Scale bars: 20 µm. (H) Western blot showing immunoprecipitation of DNMT3B and EZH2, SUZ12, EED from differentiated female ES cells. All data are presented as the mean ± SD of at least three independent experiments. \*\**P* < 0.01.

